# Emodin Combined with Multiple Low-frequency Low-intensity Ultrasound to Relieve Osteomyelitis Through Sonoantimicrobial Chemotherapy

**DOI:** 10.1101/2022.01.26.477965

**Authors:** Feng Lu, Xinhui Wu, Huiqun Hu, Zixuan He, Jiacheng Sun, Jiapeng Zhang, Xiaoting Song, Xiangang Jin, Guofu Chen

**Affiliations:** Zhejiang University School of Medicine, Hangzhou, 310009, China; Department of Orthopedic, Taizhou Hospital of Zhejiang Province, Zhejiang University, Linhai, 317000, China; Wenzhou Medical University, Wenzhou, 325035, China; Department of Orthopedic, Taizhou Hospital Affiliated to Wenzhou Medical University, Linhai, 317000, China; Department of Infectious Diseases, The Second Affiliated Hospital, Zhejiang University School of Medicine, Hangzhou, 310009, China

**Keywords:** Osteomyelitis, Emodin, Low-frequency and low-intensity ultrasound, Biofilm, Sonoantimicrobial chemotherapy

## Abstract

Treatment of osteomyelitis is still challenging as conventional antibiotic therapy is limited by the emergence of resistant strains and the formation of biofilms. Sonoantimicrobial chemotherapy (SACT) is a novel therapy of low-frequency and low-intensity ultrasound (LFLIU) combined with sonosensitizer. Therefore, in our study, a sonosensitizer named emodin (EM) was proposed to be combined with LFLIU to relieve acute osteomyelitis caused by Methicillin-resistant *Staphylococcus aureus* (MRSA) through synergistic antibacterial and anti-biofilm effects. The efficiency of different intensities of ultrasound, single (S-LFLIU, 15 min) and multiple ultrasound (M-LFLIU, 5 min every 4 h, three times) against bacteria and biofilm was compared, contributing to develop the best treatment regimen. Our results demonstrated that EM plus S-LFLIU or M-LFLIU (EM+S-LFLIU or EM+M-LFLIU) have significant synergetic bactericidal and anti-biofilm effects and EM+M-LFLIU exhibits superior performance in anti-biofilm. Furthermore, it was suggested that EM+M-LFLIU could produce a large amount of reactive oxygen species (ROS), destroy the integrity of bacterial membrane and wall, down-regulate the expression of oxidative stress, membrane wall synthesis, bacterial virulence and other related genes (*agrB, PBP3, sgtB, GMK, zwf, msrA*). *In vivo* study, micro-CT, H&E staining, ELISA assay and bacterial quantification of bone tissue indicated that EM+M-LFLIU could also relieve osteomyelitis of MRSA infection. Our work proffers an original treating bacterial osteomyelitis approach that weakens drug-resistant bacterial and suppresses biofilm formation through SACT, which may provide new prospects for clinical treatment.

## INTRODUCTION

Osteomyelitis is a difficult-to-treat bone disease often caused by a bacterial infection following a fracture or surgery. (1). It has been reported that up to 30% of high-risk open fractures will be infected, which could progress to organ infection, bone destruction, and sepsis, and is life-threatening (2, 3). The reason for the delay in curing the bacterial infection is that invasive bacteria will gradually form localized biofilms, which are organized bacterial populations encapsulated by extracellular polymeric substances (EPS) (4). Antibiotic treatment is by far the most common treatment method, but the dosage of antibiotics required to eliminate bacteria in a biofilm is 500 to 5000 times compared to the amount required for planktonic bacteria, which is beyond the human body’s ability to bear (5). In addition, antibiotic treatment often fails to completely eradicate the biofilm, leading to recurrent biofilm-associated infections (6, 7). The emergence of many drug-resistant strains also makes it imperative to find an alternative therapy to effectively remove biofilms and clear the infection (8).

Physical methods are important auxiliary methods in the struggle against bacterial drug resistance. These physical methods include the use of magnetic fields, electric fields, and ultraviolet light. Among them, low-frequency (20 kHz-1 MHz) and low-intensity (20-1000 mW/cm^2^) ultrasound (LFLIU) is a safe and promising debridement and antibacterial treatment (9, 10). The bactericidal effect of LFLIU alone is low, but combined with antibiotics can produce a synergistic bactericidal effect, whether on planktonic bacteria or bacterial biofilms (11–13). The stable cavitation of LFLIU generated numerous channels on the bacterial membrane and the extracellular matrix of biofilm that promoted the rapid entry of antibiotics into the bacteria, thus significantly increasing the effective concentration of antibiotics. Meanwhile, with the entry of more oxygen and nutrients, the sensitivity of bacteria to antibiotics can be gradually restored (14, 15). However, the use of antibiotics still retains the risk of increasing the incidence of microbial drug resistance. Therefore, developing a novel antibiotic alternative in combating the emergence of drug-resistant strains is still a great clinical challenge.

Sonoantimicrobial chemotherapy (SACT) is an emerging antimicrobial method based on a combination of sonosensitizers and LFLIU. The possible mechanisms of SACT include generating reactive oxygen species (ROS), triggering lipid oxidation, and breaking down bacterial cell membrane structures (16, 17). Besides, LFLIU can act on deep tissue microorganisms, preferentially focusing on a specific local area to activate the sonosensitizer, and ultimately exert local toxicity (18). Meanwhile, emodin (EM) is a natural anthraquinone derivative used as an alternative antimicrobial treatment. Furthermore, as a sonosensitizer, EM also has the potential to have synergistic effects with LFLIU, reportedly including anti-inflammatory, antibacterial, and anti-cancer effects (19–21).

Herein, we explored the possible synergistic effects of LFLIU and EM on planktonic bacteria and biofilm colonies. The possible influencing factors, including ultrasonic frequency (single frequency LFLIU, S-LFLIU, and multi-frequency LFLIU, M-LFLIU), ultrasonic intensity, EM concentrations, and strains were analyzed *in vitro*. The therapeutic effects of the regimens were then observed in a MRSA-infected osteomyelitis *in vivo*. The results showed that EM and LFLIU have excellent synergistic effects, whether on planktonic bacteria or biofilms. Furthermore, the combined application of LFLIU and EM was observed to effectively treat acute osteomyelitis, indicating its effective antibacterial ability *in vivo*.

## IMPORTANCE

Antibiotic therapy is the first choice for clinical treatment of osteomyelitis, but the formation of bacterial biofilm and the emergence of many drug-resistant strains also make people urgently need to find an alternative treatment to effectively eliminate the infection. Recently, LFLIU are considered to be a safe and promising method of debridement and antibacterial therapy. In this study, we found that LFLIU and EM have a significant synergistic antibacterial effect *in vivo* and *in vitro*, which may play an antibacterial role by stimulating the production of ROS, destroying the bacterial cell wall and inhibiting the expression of related genes. Our study expands the knowledge system of the antibacterial effect of EM-through combined physiotherapy. If successfully integrated into clinical practice, these methods may reduce the burden of high concentrations of drugs required for the treatment of bacterial biofilms and avoid the growing resistance of bacteria to antibiotics.

## MATERIALS AND METHODS

### Strains, Reagents and Instruments

Methicillin-sensitive *Staphylococcus aureus* (MSSA, ATCC 29213) and Methicillin-resistant *Staphylococcus aureus* (MRSA, ATCC 43300) were gained from the American Type Culture Collection. Luria-Broth (LB) and LB agar were purchased from Sangon Biotech (Shanghai, China). Distilled deionized water (ddH2O) was obtained from a Millipore purification system (USA). EM was acquired from Meilun Biotech (Dalian, China). Crystal violet was bought from Beyotime (Shanghai, China). The Live/Dead bacterial viability kit was procured from Invitrogen Molecular Probes (Eugene, OR, USA).

The LFLIU instrument (Nu-Tek, UT1021) with adjustable dual-frequency (1 MHz, 3 MHz), effective radiation area of 5.0 cm^2^ and adjustable output power of 0.1-3 W/cm^2^ was purchased from Kang Zan Medical Device. The OD values were measured using an M5 multifunctional microplate reader (SpectraMax M5, USA), fluorescence photos were taken using an SP8 LIGHTNING Confocal Microscope (Leica, Germany), and the microstructure of bacteria were photographed using a transmission electron microscope (TEM, Hitachi H-7650).

### CFU Quantification

As for antimicrobial activity *in vitro*, the bacterial suspensions were collected; diluted 10^1^, 10^2^, 10^3^, or 10^4^ times; and pointed onto LB agar plates (10 μL per point, 5 points each concentration). In addition, the 100 μL of the 10^3^ dilutions was placed onto separate LB agar plates and spread evenly for colony counting. To biofilm destruction experiment, the formed biofilms were collected, dispersed via ultrasonication for 10 min, then diluted 10^2^, 10^3^, 10^4^, or 10^5^ times to pointed onto LB agar plates, and 10^4^ dilutions were used for colony counting. To biofilm inhibition assay, the biofilm was diluted 10^0^, 10^1^, 10^2^, or 10^3^ times for pointing onto LB agar plates, and the 10^2^ dilutions were used for colony counting. As for the antibacterial effect *in vivo*, the bone homogenate was diluted 10^3^, 10^4^, 10^5^, or 10^6^ times to pointed onto LB agar plates, and 10^5^ dilutions were incubated for colony counting.

### *In Vitro* Assessment of Antimicrobial Activity

MSSA and MRSA were grown on LB medium at 37 °C for 12 h. The bacterial suspension was then diluted to 10^6^ CFU/mL for the succeeding experiments.

For the ultrasonic sterilization experiment, S-LFLIU refers to continuous irradiation for 15 min at 1 MHz frequency, and intensities of 0.1, 0.3, and 0.5 W/cm^2^ were tested, respectively. The operation frequency and intensity of M-LFLIU were the same, but the setup was irradiated for 5 min every 4 h (three times a day, tid). After corresponding irradiation, the treated bacteria were incubated at 37 °C for 6 h or 24 h to obtain the value of OD 600 to calculate their survival rate. After these initial experiments, the ultrasonic intensity of 1 W/cm^2^ induced the best antibacterial effect and was used in all succeeding experiments.

To determine the optimal concentration of EM, 50 μL double-diluted EM (1.56, 3.13, 6.25, 12.5 μg/mL) and 50 μL bacterial solution were added to 96-well plates and incubated at 37 °C for 24 h before OD 600 measurement. As for the combined antibacterial experiment, 500 μL of bacterial suspension and 500 μL double-diluted EM were added into 24-well plates, then treated with S-LFLIU (1 W/cm^2^, 15 min) and M-LFLIU (1 W/cm^2^, 5 min, tid) respectively. Then, the treated bacteria were incubated (37 °C, 24 h) and the OD 600 was measured.

We then divided the samples into six groups: control group, S-LFLIU group, M-LFLIU group, EM group, EM+S-LFLIU group, and EM+M-LFLIU group (EM: 6.25 μg/mL) for CFU counting.

### Live/dead Staining Assay

After different treatments, MRSA (10^8^ CFU/mL) were incubated (3 h, 37 °C), centrifuged and resuspended in SYTO 9/PI dye, then incubated at 37 °C for 30 min. After that, bacterial suspensions were dropped on slides to observe fluorescence images under a Confocal Microscope.

### TEM Analysis

After different treatment for 12 h (37 °C), MRSA were centrifuged and fixed overnight at 4 °C in 2.5% glutaraldehyde solution. They were then followed by a series of fixation, dehydration, permeation, embedding, sectioning and staining. Finally, the bacteria were observed under a TEM.

### Biofilm Formation and Antibiofilm Activity

To test the destruction of biofilm, MRSA were deliquated to 10^7^ CFU/mL and cultivated in 24-well plates at 37 ^o^C for 24 h to form biofilms. After that, the mature biofilms were treated for 24 h at 37 ^o^C, stained with crystal violet for 20 min and then photographed and dissolved in 95% ethanol. The damage rate in the biofilms was then calculated by comparing the OD 570 values.

After incubation for 24 h, biofilms formed on the glass slides. After a similar procedure of treatment and incubation, the formed biofilm was stained with SYTO 9 dye for 30 minutes, and its 3D structure was observed under a confocal microscope.

For the biofilm formation inhibition test, the EM and LFLIU treated MRSA (10^7^ CFU/mL) respectively at 0 h. The other experimental processes was the same as that of mature biofilm destruction test.

### ROS Detection

MRSA suspensions (10^8^ CFU/mL) were treated with corresponding treatment, incubated at 37 ^o^C for 3 h, collected and stained with DCFH-DA (Yeasen, Shanghai, China, 37 °C, 30 min). Then, bacteria were imaged under a confocal microscope. The ROS-positive rates for each sample were measured by a CytoFLEX S Flow Cytometer (Beckman Coulter, USA).

### Quantitative Real-time PCR (qRT-PCR)

The primers used in our study are listed in Table 1. After different treatment, MRSA were collected, resuspended in lysozyme solution (3 mg/mL, Meilun Biotech) and incubated at room temperature for 10 min for bacterial lysis. The RNApure Bacteria Kit (Conway Biotech, Beijing) was then used to extract the mRNA, and the mRNA was then reverse transcribed into cDNA (Takara, Beijing). qRT-PCR was then performed using a Bio-Rad CFX-96 PCR detection system (USA) and a SYBR Green I mixture (Takara, Beijing). The MRSA housekeeping gene *gyrB* was used as an internal reference for normalization of the data.

### Animal Experiments

12-week-old male C57BL/6 mice were obtained from Shanghai Slac Laboratory Animal Co., Ltd. Animal nursing and experiments were carried out in accordance with the guidelines and procedures authorized by the Animal Experimental Ethics Committee of the Taizhou Hospital of Zhejiang University.

After anesthesia, the right knee joint was shaved and sterilized, and a small incision was made on the outside of knee joint. The tibial plateau was exposed via blunt anatomy, and a single cortical bone defect of 1.5 mm in diameter was made using a dental drill. MRSA (1×10^8^ CFU/mL, 10 μL) were inoculated into the marrow of the right bone defect, and then the muscle fascia and skin were sutured. After the operation (day 0), the animals were equally divided into the following groups: (1) blank group,; (2) control group; (3) M-LFLIU group; (4) EM group; (5) EM+M-LFLIU; The treatment method was performed according to the following procedure: remove the sutures from the surgical incision of the mouse, separate and expose the tibial bone hole, and use M-LFLIU (1 W/cm^2^, 5 min, tid) and 10 mL of EM solution (40 μg/mL) to wash the wound and bone hole (fluid flow rate: 10 mL/min) according to the treatment group, then apply a local EM wet compress, and wrap the wound with a sterile dressing. During this period, the body temperature and bodyweight of mice were measured every other day. After 7 days, the right leg and bone samples of mice were collected for photographing and bacterial quantification

### Micro-CT Analysis

A high-resolution micro-CT (μCT-100, SCANCO Medical AG, Switzerland) was used to explore the tibia and knee joint fixation. The scanning scheme used was as follows: equal resolution 20 μm, 70 kV, 200 μA, and exposure time 300 ms. After reconstruction, the degree of the destruction of these structures in each group was observed and compared. In each sample, both the proportion of total porosity and BV/TV (bone volume/ total volume) were assessed by using Scanco software (SCANCO Medical AG, Switzerland; Evaluation V6.5-3).

### ELISA

The expression level of IL-6, IL-1α, and IL-1β in bone homogenate were measured by ELISA kits provided by AMEKO (Hangzhou, China). All the procedures were performed compliant with the manufacturer’s instructions.

### *In Vivo* Toxicity Assay

Two weeks after the treatment, blood routine and blood biochemical were conducted to explore the toxicity of EM+M-LFLIU *in vivo*. The heart, liver, spleen, lung, and kidneys were stained with H&E for the determination of toxicological histology.

### Statistical Analysis

All the data in this study were statistically analyzed by Student’s t-test (unpaired, two-tailed). The value was expressed as mean ± standard mean error (SEM), and P < 0.05 was considered statistically significant (* P < 0.05, ** P < 0.01).

## RESULTS

### The Antibacterial Efficiency of EM and LFLIU

The survival rate of MSSA was evaluated using the OD 600 value after incubating for 6 h or 24 h, respectively. When the intensity was increased to 1 W/cm^2^, the survival rate decreased significantly right after 6 h, but showed no obvious difference at 24 h (**Figure 1A**). Similar results can be seen after M-LFLIU (**Figure 1B**) treatment, indicating that the antibacterial effect of LFLIU treatment is temporary.

**FIGURE 1.**
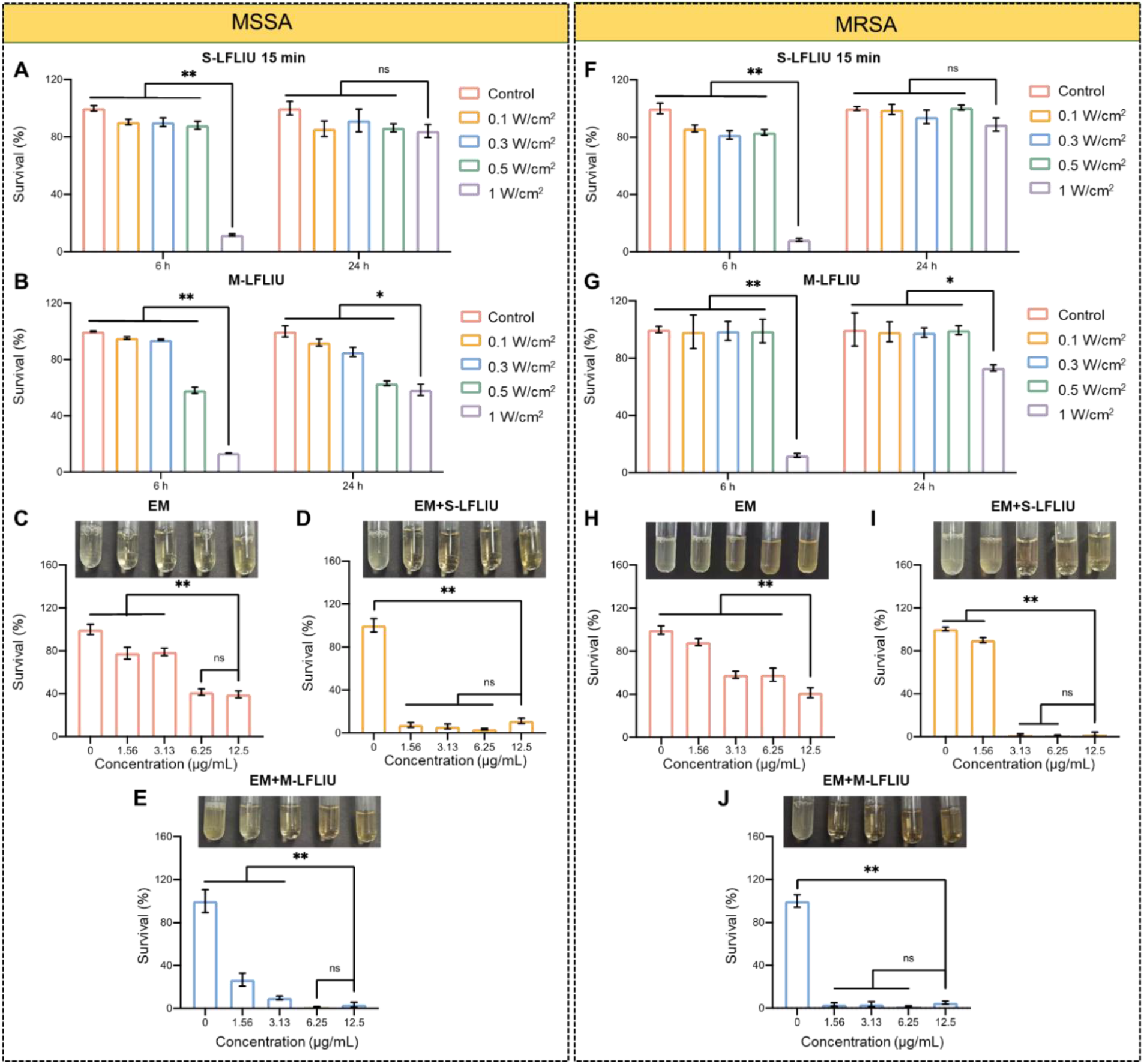
**(A-B)** The survival rate of MSSA treated with S-LFLIU or M-LFLIU at different frequencies for 6 hours and 24 hours, respectively **(C-E)** The survival rate of MSSA treated with different concentration of EM, EM+S-LFLIU or EM+M-LFLIU respectively at 37 °C for 24 hours (S-LFLIU:15 min, M-LFLIU: 5 min every 4 h, three times). **(F-G)** The survival rate of MRSA treated with S-LFLIU or M-LFLIU at different frequencies for 6 hours and 24 hours, respectively. **(H-J)** The survival rate of MRSA treated with different concentration of EM, EM+S-LFLIU or EM+M-LFLIU, respectively.

We further studied the antimicrobial activity of EM alone or combined with S-LFLIU or M-LFLIU on MSSA. When the concentration of EM was increased, the survival rate of MSSA decreased. However, the survival rate was still as high as ~39.4% even when the concentration was increased to 12.5 μg/mL (**Figure 1C**). The survival rate of the bacteria was obviously lower in EM combined with S-LFLIU or M-LFLIU, which was less than 10%, with the bacterial suspension appearing clear after incubation for 24 h (**Figure 1D, E**). Similar results were seen in parallel experiments on MRSA (**Figure 1F-J**).

To further investigate the antibacterial activity of EM plus S-LFLIU or M-LFLIU, we observed the viable bacterial counts on LB agar plates (**Figure 2A, B**). Viable bacterial counts decreased in the EM group to some degree, and relatively few living colonies were found in both the EM+S-LFLIU and EM+M-LFLIU groups.

**FIGURE 2.**
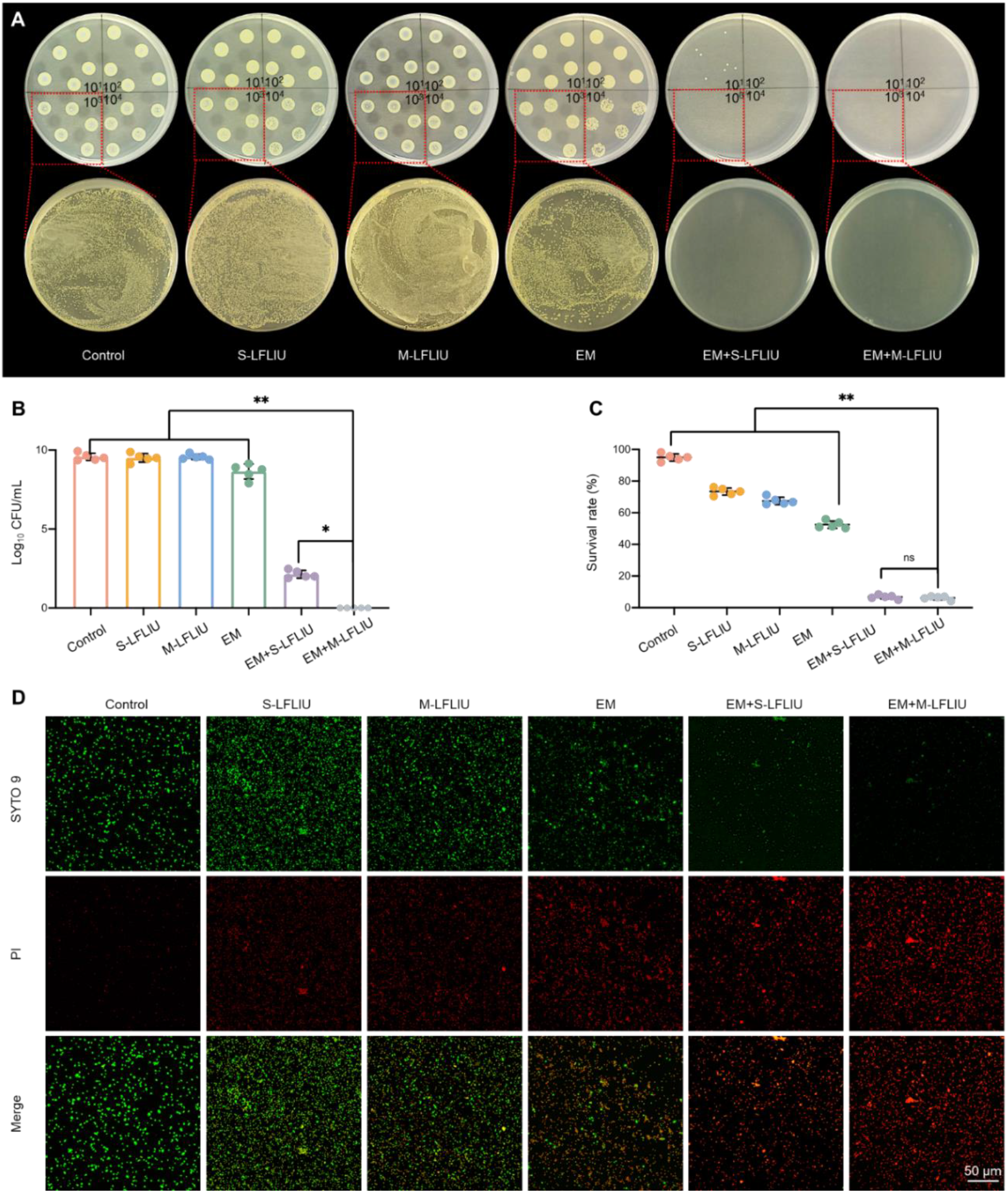
**(A)** Photos of the growth of MRSA on the LB agar plates treated with blank LB medium, S-LFLIU, M-LFLIU, EM, EM+S-LFLIU or EM+M-LFLIU respectively at 37 °C for 24 hours (S-LFLIU:15 min, M-LFLIU: 5 min every 4 h, three times), (top: diluted 10^1^, 10^2^, 10^3^, 10^4^ times, bottom: diluted 10^3^ times). **(B)** Quantitative analysis of the number of MRSA after different treatment. **(C-D)** Quantitative and qualitative analysis and live/dead staining images of MRSA after different treatment.

The live/dead staining of MRSA clearly showed the proportion of living bacteria to the total bacterial community was decreased in the EM group, and a greater decrease in the EM+S-LFLIU and EM+M-LFLIU groups. (**Figure 2C, D**). These results indicated a significant synergistic effect of EM and LFLIU, and no significant difference was found between S-LFLIU and M-LFLIU in terms of enhancing the bactericidal effect of EM.

### Synergistic Activity EM plus LFLIU Against MRSA Biofilm

Considering the significant bactericidal activity of EM plus LFLIU on planktonic MRSA, we next studied its effect on MRSA biofilms. As shown in **Figure 3A and B**, the stained biofilms were slightly thinner in the M-LFLIU and EM groups and much thinner in the EM+S-LFLIU and EM+M-LFLIU groups. The inhibition rates of the above groups were 7.0%, 30.9%, 56.8%, 75.6%, and 98.5%, respectively. Interestingly, after the same treatment, the formed biofilm could also be damaged to different degrees, with destruction rates of 6.1%, 40.2%, 69.9%, 83.6%, and 95.5%, respectively (**Figure 3D, E**). **Figure 3C and F** showed the 3D structure of biofilm after corresponding early- and late-stage treatments. It was observed that the EM+M-LFLIU group had the weakest green signal and thinnest thickness, which implies that the strongest anti-biofilm ability belongs to EM+M-LFLIU.

**FIGURE 3.**
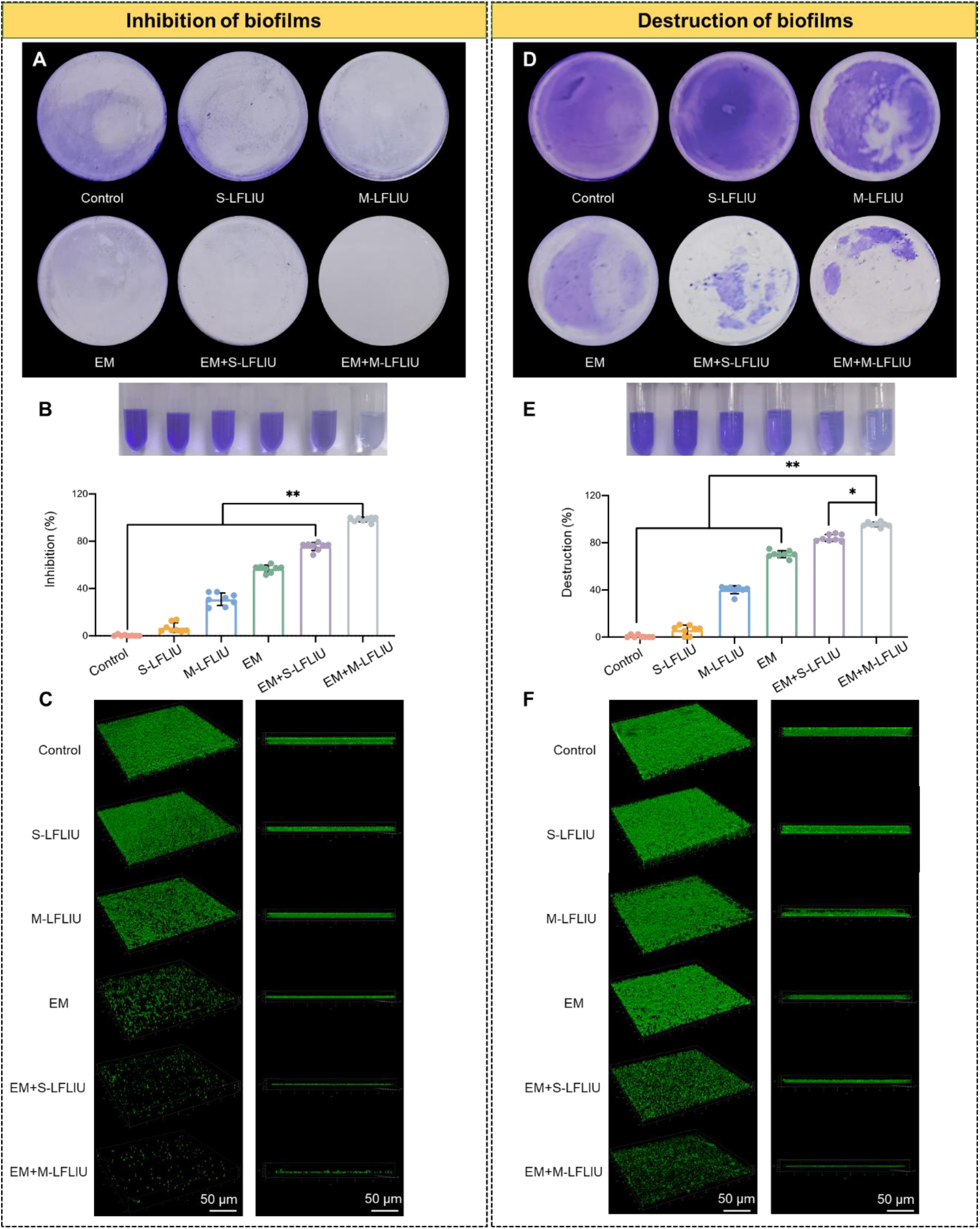
**(A)** Crystal violet staining photos of immature MRSA biofilms (treated at 0 h) after different treatment. **(B)** Inhibition rate of MRSA biofilm formation calculated by quantitative analysis of crystal violet staining (upper: photo of crystal violet stained biofilm dissolved in ethanol) **(C)** 3D fluorescence image of immature MRSA biofilm took by laser confocal microscope (left: top view right: side view). **(D)** Crystal violet staining photos of mature MRSA biofilms (treated at 24 h) after different treatment. **(E)** Destruction rate of mature MRSA biofilm calculated by quantitative analysis of crystal violet staining (upper: photo of crystal violet stained biofilm dissolved in ethanol). **(F)** 3D fluorescence image of mature MRSA biofilm took by laser confocal microscope (left: top view right: side view)

Bacterial plate counting was performed to assess the load of live MRSA in treated biofilms. The photo of colonies grown on the plates are depicted in **Figure 4A and 4B**, and the quantitative statistics of colony numbers are presented in **Figure 4C and 4D**. Compared with the blank LB medium treatment, the number of colonies was evidently lessened in the EM groups, with an even greater decrease noted in the EM+S-LFLIU and EM+M-LFLIU groups whether the biofilm was treated at the early stage or the late stage. Moreover, nearly no bacteria were found after treatment with EM+M-LFLIU. Taken together, these results support that EM plus LFLIU showed synergistic anti-MRSA biofilm activity and that EM+M-LFLIU performed better than EM+S-LFLIU in achieving this.

**FIGURE 4.**
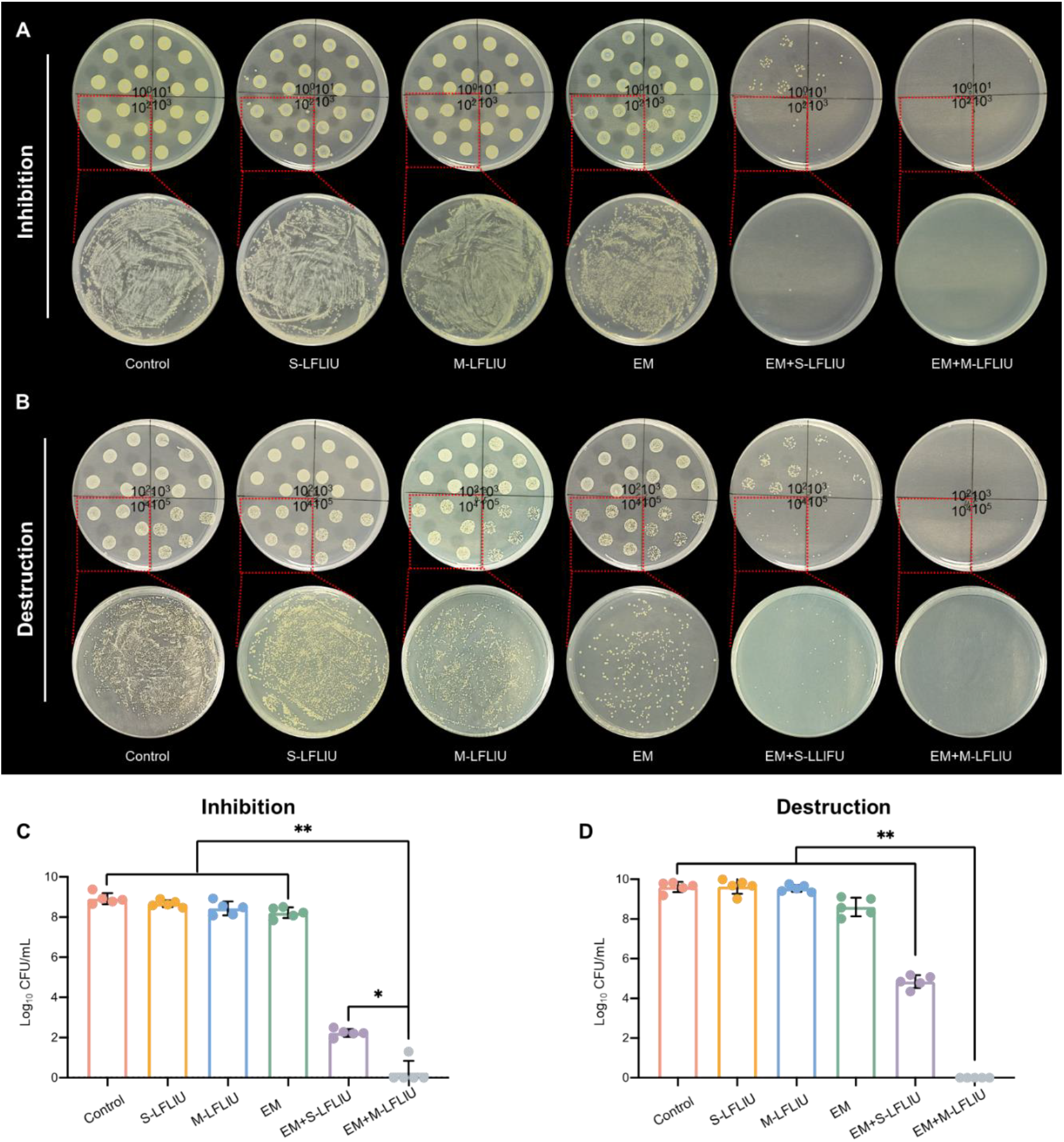
**(A)** The picture of the growth of viable MRSA in the immature biofilm on the LB agar plates after the corresponding treatment (top: diluted 10^0^, 10^1^, 10^2^, 10^3^ times, bottom: diluted 10^2^ times). **(B)** The picture of the growth of viable MRSA in the mature biofilm on the LB agar plates after the corresponding treatment (top: diluted 10^2^, 10^3^, 10^4^, 10^5^ times, bottom: diluted 10^4^ times). **(C)** Quantitative analysis of the number of MRSA colonies in the immature biofilm after corresponding treatment. **(D)** Quantitative analysis of the number of MRSA colonies in the mature biofilm after corresponding treatment.

### Potential Antibacterial Mechanism of EM plus LFLIU

In order to further investigate its potential antibacterial mechanism, the production of ROS was detected. As revealed in **Figure 5A** and **5B**, the strongest ROS signal was observed when bacteria were treated with EM plus M-LFLIU, while a weak ROS signal in EM alone, and almost none in the control and M-LFLIU-only groups. Similarly, the flow cytometry results (Figure 5C) suggest that a large amount of ROS was produced in EM+M-LFLIU group, participating in the sterilization process.

**FIGURE 5.**
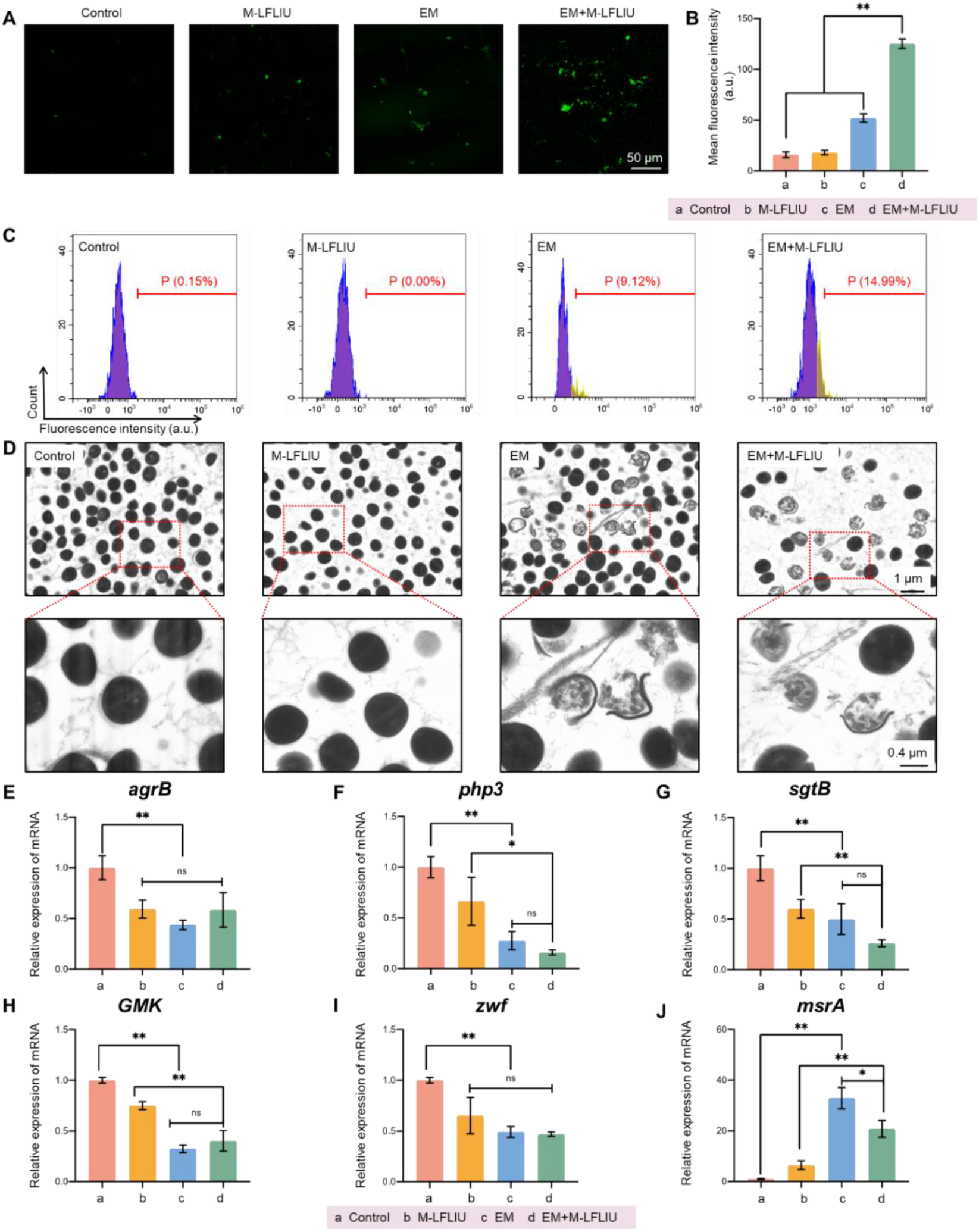
**(A-B)** Fluorescence image and quantitative analysis of ROS generated after treated with blank LB medium, LFLIU, EM and EM+LFLIU. **(C)** Positive rate of ROS production detected by Flow Cytometer in different groups. **(D)** TEM images of MRSA in different group (Below are enlarged views of the upper). **(E-J)** Relative mRMA expression of *agrB, PBP3, sgtB, GMK, zwf, msrA* measuered by qRT-PCR in different groups.

As for the morphological changes, the treated bacteria were photographed under a TEM (**Figure 5D**). It was observed MRSA in the control and M-LFLIU groups remained intact. However, the bacteria treated with EM and EM+M-LFLIU lost bacterial wall integrity and their cell contents leaked to a large extent. Additionally, the EM+M-LFLIU group showed fewer bacteria and a larger proportion of bacteria presenting with bacterial membrane rupture.

Subsequently, we have conducted focused research and screening on the expression of some genes related to bacterial growth and synthesis, bacterial virulence, and oxidative stress. First, *agrB* is responsible for encoding a co-gene regulatory protein B located in the plasma membrane, which is associated with vancomycin sensitivity and with the virulence of MSSA (22–25). In regulating bacterial growth and synthesis, penicillin-binding protein 3 (*PBP3*) is involved in the final stage of peptidoglycan biosynthesis, thus working in the formation of the cell wall. Meanwhile, monofunctional glycosyltransferase (MGT) has been shown to participate in the extension of the peptidoglycan chain, while *PBP3* and *sgtB* are the genes responsible for encoding the above two proteins respectively (23, 26, 27). *GMK*, which encodes cytoplasmic guanosine kinase, is involved in the synthesis of nucleotide precursors and indirect regulation of DNA and RNA synthesis. Inhibition of *GMK* can affect bacterial growth; therefore, it is a potential target for new antimicrobial agents against MSSA (23, 28). In terms of bacterial regulation of oxidative stress, we found that *zwf* and *msrA* take part in protecting cells from oxidative stress (23). The upregulation of *zwf* has been proven to be a response to oxidative stress, heat stress, and even virulence (29–31). On the other hand, *msr* encodes methionine-S-sulfoxide reductase, the downregulation of which may affect the resistance to oxidative stress (32, 33). Then, our experimental results showed that M-LFLIU and EM could inhibit their expression in terms of bacterial virulence and oxidative stress-related genes, but they have no obvious synergistic effect. In bacterial growth and synthesis-related genes, EM inhibits the expression of cell wall synthesis-related genes, and M-LFLIU appears to play a synergistic effect (**Figure 5E-J**).

In summary, EM+M-LFLIU appear to exert antibacterial effects on MRSA by producing ROS and destroying the bacterial cell wall while reducing the expression of genes related to bacterial cell wall synthesis. Besides, EM+M-LFLIU may weaken the virulence of MRSA, reduce its toxic and side effects on the host, weaken the resistance of MRSA to oxidative stress and strengthen the bactericidal effect of ROS.

### *In Vivo* Effect of EM+M-LFLIU on Osteomyelitis Caused by MRSA

Encouraged by the results *in vitro*, the *in vivo* effect of EM+M-LFLIU on osteomyelitis caused by MRSA was also investigated. A MRSA-infected osteomyelitis model was established to evaluate the efficacy *in vivo* (**Figure 6A**). On day 7, the legs and bone tissue of each group are shown in **Figure 6B**. Contrast to the blank group, the right legs of the no treating and the M-LFLIU group were obviously enlarged, with a greater number of small pustules scattered on the surface of muscle tissue. There was slightly less swelling and lower volume of suppurative tissue in the EM group, inconspicuous swelling and no scattered pustules in the EM+M-LFLIU group.

**FIGURE 6.**
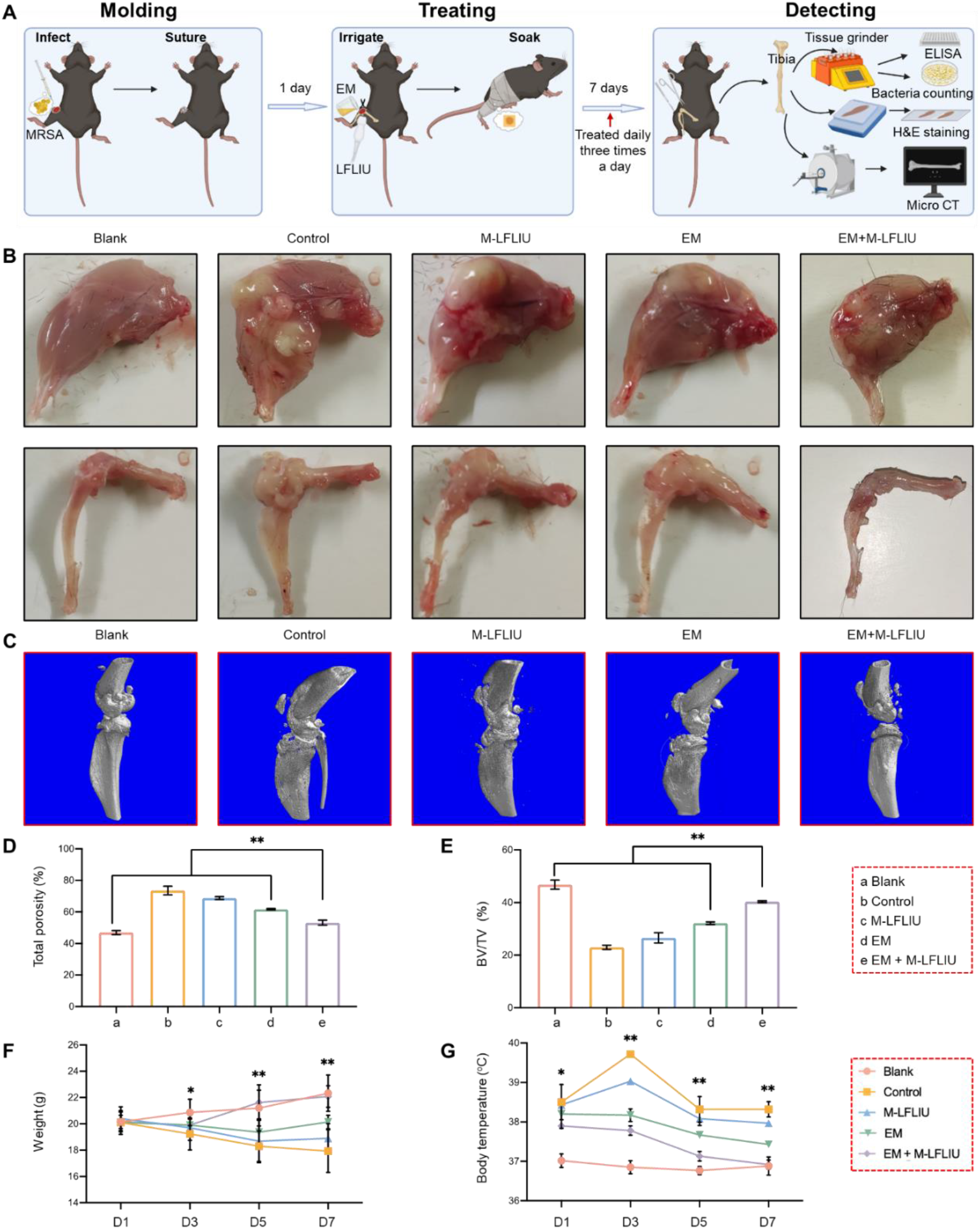
**(A)** Schematic diagram of the animal experiment process. **(B)** The photos of mouse legs (top: legs with muscle and bone, bottom: legs with bone alone). **(C)** Micro-CT 3D reconstruction of the leg bones in different groups. **(D)** Total porosity of tibia in different groups. **(E)** BV/TV of tibia in different groups. **(F-G)** Mouse weight and body temperature recorded every other day in different groups.

After the muscle and soft tissue were removed, the appearance of bone tissue in different groups were observed. Compared with the blank group, the bone destruction of the knee joint in the control, M-LFLIU, and EM group was more severe, the tibia seemed to be swollen, and suppurative tissue could be seen in the bone marrow cavity. In the EM+M-LFLIU group, bone destruction was less pronounced, the volume of the tibia was not significantly increased, and the suppurative tissue in the bone marrow cavity was not obvious. The micro-CT images obtained further corroborates the above findings (**Figure 6C**). The results showed that the BV/TV in the control group was the lowest and increased after M-LFLIU or EM treatment, while the EM+M-LFLIU group was the closest to that in the blank group, and the total porosity showed the opposite trend, which was consistent with the micro-CT images (**Figure 6D, E**).

In addition, over the course of treatment, we recorded the body temperature and body weight of each group every other day to determine the basic condition of the mice. Contrary to the body weight gain of mice in the blank group, the bodyweight of untreated mice continuously decreased. The mice in the EM+M-LFLIU group had very little weight loss and started to increase steadily on the 3rd day after treatment, and even approaching that of blank group at the end of the treatment period. In terms of body temperature, the mice in the no treating group and the M-LFLIU group reached highest level on the 3rd day after treatment and then decreased to approximately 38 °C, while in the EM and EM+M-LFLIU groups reached above 38 °C on the first day of treatment and then gradually decreased to 37.9 ^o^C and 37.4 ^o^C, respectively. In particular, there was no obviously difference between the EM+M-LFLIU and the blank group on the 7th day after treatment (**Figure 6F, G**).

### Antibacterial and Anti-inflammatory Effects of EM+M-LFLIU *In Vivo*

The antibacterial effects were evaluated via bacterial plate counting and ELISA. The pictures of colonies on the LB agar plates and the quantitative statistics of the colony count numbers are shown in **Figure 7A and B**. The ELISA results of inflammatory factors in bone tissue are shown in **Figure 7C-E**.

**FIGURE 7.**
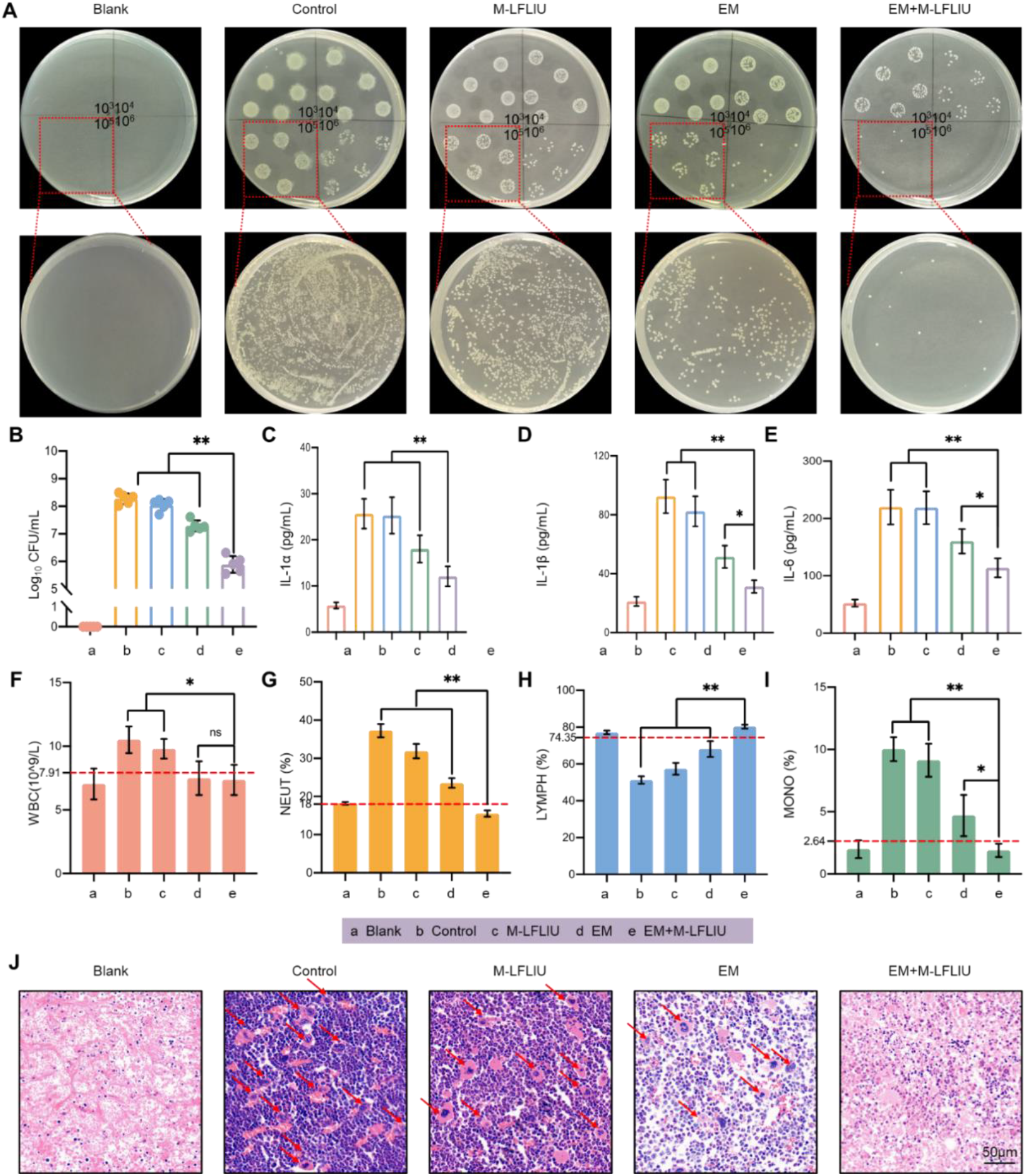
The picture of the growth of bacteria in the homogenate of bone tissue on the LB agar plates 7 days after the corresponding treatment (top: diluted 10^3^, 10^4^, 10^5^, 10^6^ times, bottom: diluted 10^5^ times). **(B)** Quantitative analysis of the number of viable bacterial colonies in homogenate of bone tissue 7 days after corresponding treatment. **(C-E)** The expression level of inflammatory factors (IL-1α, IL-1β, IL-6) tested by ELISA kit. **(F-I)** The number of white blood cells, the proportion of neutrophils, the proportion of lymphocytes, and the proportion of monocytes in the blood routine test 7 days after corresponding treatment. The red dotted line represents the normal value. **(J)** H&E staining images of bone (tibia) 7 days after corresponding treatment (red arrows represent typical inflammatory cells).

Contrast with the untreated group, the number of colonies and the expression levels of IL-1α, IL-1β and IL-6 were slightly lower in the M-LFLIU group, and evidently reduced in the EM group and EM+M-LFLIU group. Moreover, EM+M-LFLIU has the least number of colonies and the lowest level of inflammatory factors. Consistent with these findings, H&E staining of bone tissue sections also showed that the control group had a large number of inflammatory cells, suggesting severe inflammation and the EM+M-LFLIU inflammation in the group was greatly alleviated (**Figure 7F**).

Finally, a complete blood count was performed to estimate the degree of inflammation in the mice. The blood test results indicated that the inflammation of the EM group was suppressed to a certain extent, while the inflammation of the EM+M-LFLIU group mice was effectively controlled (**Figure 7G-J**). As the degree of inflammation in mice intensifies, the number of leukocytes, and the ratio of neutrophils and monocytes will increase significantly, while the ratio of lymphocytes will decrease. Therefore, we suggest that EM+M-LFLIU has excellent potential for effectively killing bacteria and decreasing the levels of inflammatory markers.

### Pilot Toxicity Testing *In Vivo*

We then tested whether treatment with EM+M-LFLIU has a toxic effect on mice. Histological structures of the heart, liver, spleen, lung and kidney (**Figure 8A**), the complete blood count, and the hepatic/renal function indices (**Figure 8B-I**) all indicated that EM+M-LFLIU did not induce any significant alterations in mice, indicating the excellent biosafety at the therapeutically efficacious dose *in vivo*.

**FIGURE 8.**
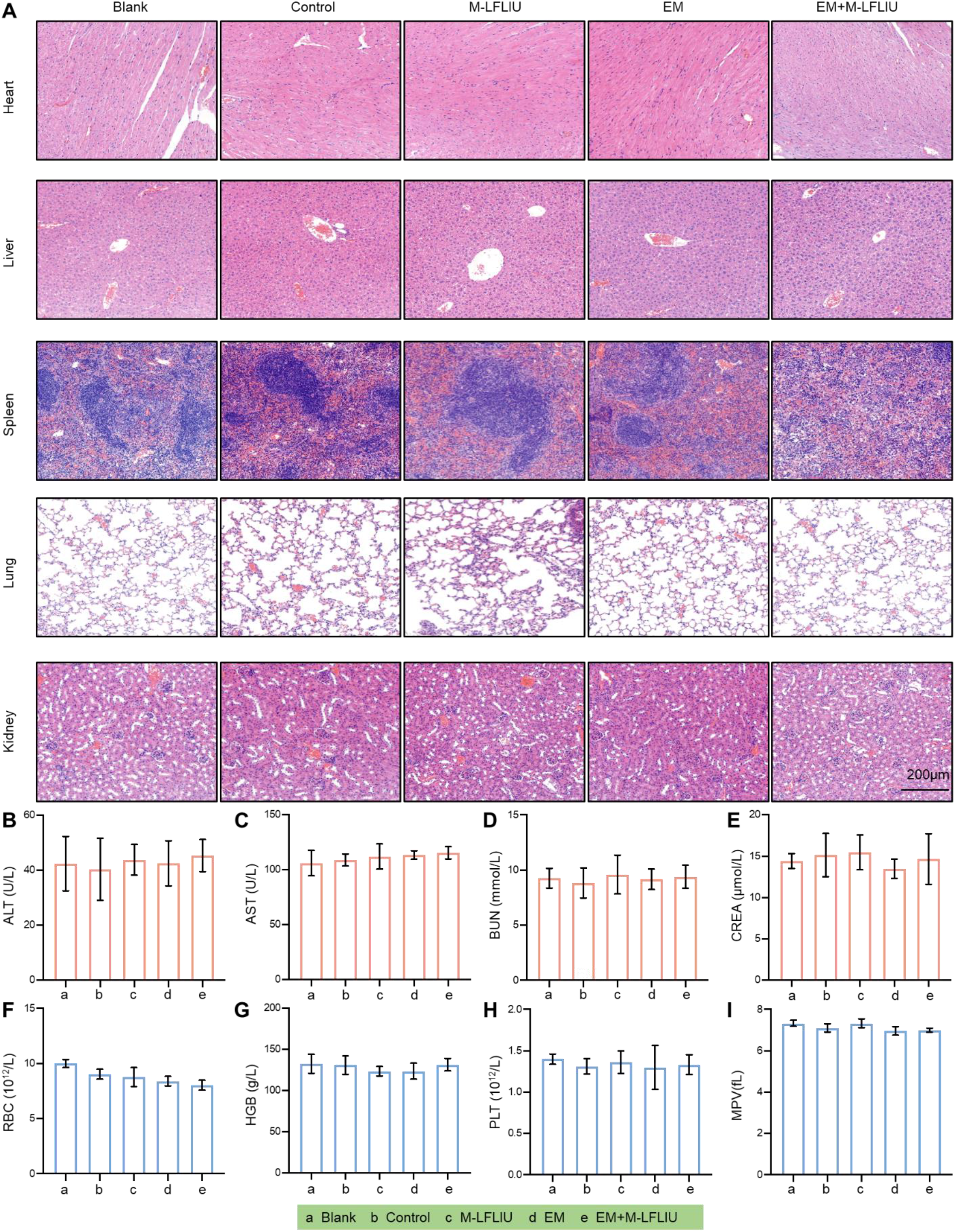
**(A)** H&E staining images of major organ (heart, liver, spleen, lung, kidney) two weeks after the completion of the treatment in different groups. **(B-E)** Blood biochemistry after two weeks of treatment in different groups (ALT, AST, BUN, CREA). **(F-I)** Blood routine after two weeks of treatment in different groups (RBA, HGB, PLT, MPV).

## DISCUSSION

Traumatic osteomyelitis, which often develops a MSSA infection, is the most common type of osteomyelitis (34). Vancomycin is recognized as an effective drug for the clinical treatment of acute osteomyelitis (35). However, the formation of bacterial biofilms can reduce the effectiveness of antibiotics. With the increasing prevalence of antibiotic resistance in bacteria (36), developing new antibacterial drugs and alternative therapies is imperative. EM is a drug with significant anti-inflammatory and antibacterial activity (37, 38), thus it has potential to replace antibiotics as an effective drug for treating acute osteomyelitis. Previous studies have reported the antibacterial effect of ultrasound in food packaging (39), and LFLIU has also been reported to stimulate ROS production, which provides a specific repressive effect on the growth of suspended bacteria and bacterial biofilms (12, 13, 40). In this study, it found that M-LFLIU combined with EM had significant synergistic antibacterial effects *in vitro* and *in vivo*, providing a promising possibility for alternative clinical treatments of acute osteomyelitis.

We first explored the antibacterial effect of LFLIU of different intensities and frequencies on MSSA and MRSA. The experimental results suggest that the antibacterial effect of a single application of LFLIU or EM is not ideal, and is difficult to achieve the desired effect. After combined treatment with EM and LFLIU, even low concentrations of EM can have a satisfactory antibacterial effect. This reminds us that the combined use of different measures can achieve the effect of one plus one greater than two, and can effectively avoid the side effects caused by the excessive degree of a single measure. Our experimental results verify and expand the previous hypothesis that LFLIU combined with EM has a synergistic antibacterial effect.

The formation of bacterial biofilm is widely regarded as an important part of the bacterial infection process and a major challenge in treating an infection (41). Mature bacterial biofilm can effectively resist the antibacterial effect of antibiotics. Therefore, we studied the effects of EM and LFLIU on the growth of suspended bacteria and the formation and destruction of bacterial biofilms. It is seen that EM+LFLIU, especially EM+M-LFLIU, showed a clear effect in the retardation of bacterial growth and bacterial biofilm formation, and increased the destruction of the biofilms. From this, we concluded that the antibacterial effect of M-LFLIU combined with EM is an effective and promising plan for clinical treatment. It is expected to solve the problems that antibiotics are difficult to solve the biofilm barrier and the production of drug-resistant bacteria.

After observing the remarkable results, we began to explore the antibacterial mechanism of EM+M-LFLIU. The TEM results showed that EM could effectively cause the rupture of the bacterial cell wall. The proportion of bacteria with cell wall rupture increased after M-LFLIU. Since EM has broad-spectrum effects, including anti-inflammatory, antioxidant, and antibacterial (42), we tested ROS production in our system. Confocal probe and flow cytometry showed that EM could stimulate ROS production and exert its antibacterial effect and that M-LFLIU could amplify this effect. Finally, we focused on the expression of bacterial genes related to growth synthesis, virulence, and oxidative stress defense, such as: *agrB, php3, sgtB, GMK, zwf and msrA*. The results showed that EM+M-LFLIU may weaken the virulence of MRSA, reduce its toxic and side effects on the host, weaken the resistance of MRSA to oxidative stress and strengthen the bactericidal effect of ROS. Thus, these imply a potential antibacterial mechanism of EM and provide some directions for future research of the synergistic effect of EM+LFLIU. In summary, LFLIU combined with EM appear to exert antibacterial effects on MRSA by producing ROS and destroying the bacterial cell wall while reducing the expression of genes related to bacterial cell wall synthesis.

To determine the effectiveness of EM with M-LFLIU *in vivo*, we made a mouse model to test the therapeutic effect on acute osteomyelitis by observing the bone and the surrounding tissue abscess, cortical bone loss, basic vital signs, and related inflammatory indexes. Compared with the traditional administration methods such as intragastric administration or intraperitoneal injection, our animal model was rinsed with aseptic saline containing emodin and given local wet compress, which effectively increased the local effective drug concentration, avoided the possibility of toxic side effects caused by systemic administration of excessive emodin, better combined with ultrasound to form local targeting, and was more in line with clinical treatment. As expected, the symptoms of osteomyelitis in mice treated with M-LFLIU combined with EM were significantly alleviated. The degree of bone destruction was mild, and the inflammatory indexes and vital signs recovered rapidly compared to the other groups. Finally, the blood biochemistry and visceral tissue section H&E staining indicated that EM had no obvious toxic side effects. In total, this implies that EM+LFLIU is a clinically viable treatment for acute osteomyelitis.

In summary, our study expands the knowledge of the antibacterial effect of EM–through combined physiotherapy. If successfully integrated into clinical practice, these methods may reduce the burden of high concentrations of drugs needed to treat bacterial biofilms and avoid the growing resistance of bacteria to antibiotics. We have shown that this method provides a meaningful possibility for treating osteomyelitis caused by MRSA. Future research may show that it is useful in treating infections and conditions caused by other antibiotic-resistant bacteria.

## DATE AVAILABILITY STATEMENT

The raw data supporting the conclusions of this article will be made available by the authors, without undue reservation.

## ETHICS STATEMENT

The study was conducted according to the guidelines of the Declaration of Helsinki, and approved by the Institutional Animal Ethics Committee of Taizhou Hospital. (protocol code: tzy-2021164)

## AUTHOR CONTRIBUTION

GC and FL conceived the study. FL, HH contributed to the acquisition and interpretation of data. FL and XW did the data cleaning, statistical analysis and model building with the assistance from ZH, JS, JZ, XS and XJ. FL and HH drafted the manuscript. All authors contributed to critical revision of the manuscript for important in- tellectual content. All authors critically reviewed the manuscript for important intellectual con- tent and gave final approval for the version to be published.

